# Performance Characteristics of Zeno Trap Scanning DIA for Sensitive and Quantitative Proteomics at High Throughput

**DOI:** 10.1101/2025.05.06.652368

**Authors:** Ludwig R. Sinn, Ziyue Wang, Claudia P. Alvarez, Anjali Chelur, Ihor Batruch, Patrick Pribil, Daniela Ludwig, Stephen Tate, Jose Castro-Perez, Christoph B. Messner, Vadim Demichev, Markus Ralser

## Abstract

Proteomic experiments, particularly those addressing dynamic proteome properties, time series, or genetic diversity, require the analysis of large sample numbers. Despite significant advancements in proteomic technologies in recent years, further improvements are needed to accelerate measurement and enhance proteome coverage and quantitative performance. Previously, we demonstrated that incorporating a scanning MS2 dimension into data-independent acquisition methods (Scanning SWATH, or more generally scanning DIA) but also ion trapping, improves analytical depth and quantitative performance, especially in proteomic methods using fast chromatography. Here, we evaluate the scanning DIA approach combined with ion trapping via the Zeno trap in a method termed ZT Scan DIA, using a prototype Zeno trap Q-TOF instrument (SCIEX). Applying this method to established proteome standards across various analytical setups, enabling intermediate to high sample throughput, we observed a 30–40% increase in identified precursors. This enhancement extended to overall protein identification and precise quantification. Furthermore, ZT Scan DIA effectively eliminated quantitative bias, as demonstrated by its ability to deconvolute proteomes in multi-species mixtures. We propose that ZT Scan DIA can be used for a broad range of applications in proteomics, particularly in studies requiring high quantitative precision with low sample input and high-throughput workflows.

**Significance Statement:** The advent of faster DIA proteomics methods paved the way for investigating increasingly large sample series, including patient cohorts and strain collections containing thousands of samples. Yet, recent improvements in DIA methods still entailed compromises between analytical sensitivity and selectivity. The presented combination of a scanning quadrupole with fast ion trapping in a Zeno Trap, coined ZT Scan DIA increases quantitative precision and accuracy in fast proteomics experiments. These features of ZT Scan DIA may benefit applications that deal with low input samples and high-throughput proteomic workflows in biomedical cohort studies and systems biology.

## 1 Introduction

LC-MS-based proteomics expanded from predominantly protein identification experiments in basic science to increasingly quantitative and translational applications, including large-scale system biology or plasma proteomic studies. These transitions were facilitated by numerous technological improvements that increased measurement precision and throughput (Aebersold & Mann, 2016; Messner et al., 2023). These enhancements include increasingly automated sample preparation, improved data processing software, and better instruments to perform sophisticated LC-MS analyses that are faster and more sensitive (Bian et al., 2020; Cardozo et al., 2020; Demichev et al., 2020; Fu et al., 2018; Hendricks et al., 2024; Kong et al., 2017; Szyrwiel et al., 2024; Vowinckel et al., 2018).

Mostly due to the promise to minimize stochastic elements and to improve quantification performance, data-independent acquisition (DIA) schemes gained in popularity, especially for experiments that involve a high number of samples (Deutsch et al., 2021; Fröhlich et al., 2022; Gonçalves et al., 2022). One of the first commercially successful DIA methods was SWATH-MS, implemented on the Triple TOF 5600 instrument (a QTOF, quadrupole time-of-flight, SCIEX) (Gillet et al., 2012). When coupled with capillary flow chromatography, SWATH-MS could generate precise proteomic measurements over hundreds of samples, in runtimes of 30 minutes or less per sample (Vowinckel et al., 2018) (Wang et al., 2022) (Fig. 1a,d). The second generation of SWATH-MS allowed for the window sizes to be modulated depending upon the complexity of the sample. Although not improving the overall cycle time, this variable window SWATH allowed for improved selectivity and performance over the original methodology (Zhang et al., 2015). A third generation of the SWATH method benefitted from the integration of a novel linear ion trap (“Zeno trap”), which increased sensitivity and improved proteomic depth (Zeno SWATH MS) (Wang et al., 2022) (Fig. 1b,e). In addition, the Zeno SWATH method reduced the demand for sample amounts, mitigating matrix effects and enhancing the longevity of ion optics and the detection system. Although sample throughput could be increased by allowing shorter ion accumulation times thanks to overall improved sensitivity, rapid advances in chromatography (Bache et al., 2018; Sun et al., 2023) pushed a need for faster cycle times. To overcome this limit, we have previously presented the Scanning SWATH approach (scanning DIA), in which the windowed acquisition of SWATH methods was replaced with a scanning quadrupole mode (Messner et al., 2021). In this DIA mode, the Q1 quadrupole is operated in a mode identical to a Q1 scan on a Triple Quadrupole instrument. Isolated precursors are transmitted to the collision cell as the subsequent fragment ions are observed in multiple subsequent TOF MS2 pulses (Fig. 1c,d). Consequently, every detected ion is characterized by four dimensions: intensity, *m/z*, chromatographic retention time, and the position of the Q1 window at which this ion has been detected. Analysis of the observed fragment ion intensity profiles across the varying Q1 isolation window positions enables linking individual MS/MS signals to a particular peptide precursor ion *m/z*. Furthermore, the Q1 dimension aids in identifying and excluding signal interferences that arise in the presence of contaminants, ion adducts, or losses, as well as modified peptide derivatives at a close *m/z*. Exploiting the Q1 scan dimension to improve precursor identification and quantification in our DIA-MS software DIA-NN (Demichev et al., 2020), we demonstrated ultra-high throughput proteomic experiments in combination with 800µl min^-1^ high-flow rate liquid chromatography and chromatographic gradients that were as fast as 30 seconds (Messner et al., 2021).

**Fig. 1:**
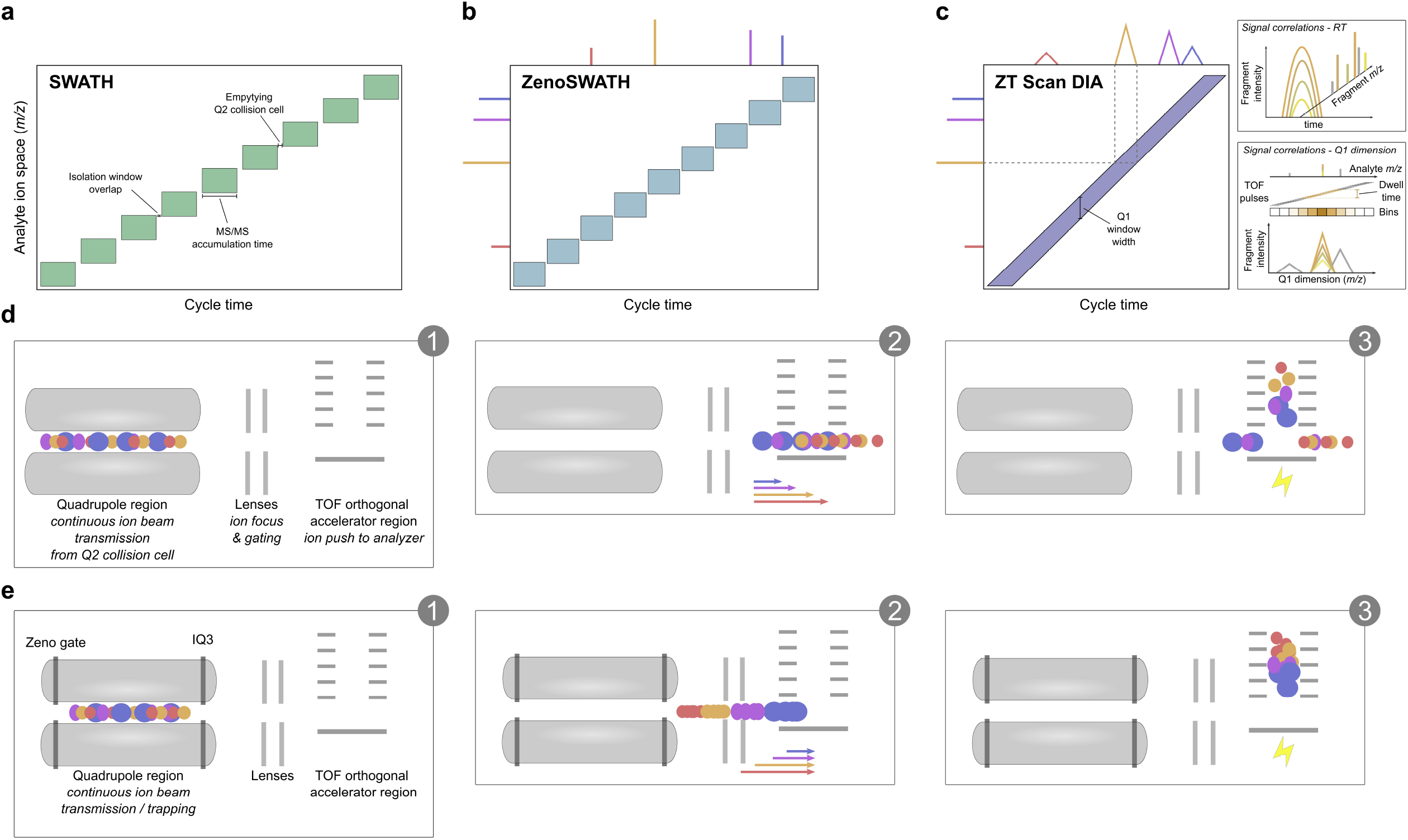
Scheme on SWATH / Zeno SWATH / ZT Scan DIA modes of operation. Comparison of SWATH, Zeno SWATH, and ZT Scan DIA regarding ion isolation, fragment ion handling, and ZT Scan DIA data structure. Panel (a) shows the SWATH principle, in which ions are isolated and fragmented in sequential windows, each with a small overlap in the *m/z* dimension. Due to time expenses when isolating, fragmenting, and fragment ion ejection from the Q2 collision cell, the time for a sequence to complete a cycle (“cycle time”) is comparably long. In panel (b), Zeno SWATH allows shorter cycle times since the time demand for emptying Q2, and ion losses in general, are reduced due to the use of the Zeno trap for trapping ions (see panel e). Still, fragment ions from a given peptide precursor will be observed per individual isolation window. Panel (c) showcases the ZT Scan DIA mode in which the Q1 quadrupole rapidly scans the precursor ion space, thus enabling to leverage fragment ion signal correlations in the retention and *m/z* x time domains (panel (c) right). Panel (d) displays ion handling in the Quadrupole and TOF pulser regions as in SWATH and Scanning DIA. A continuous ion beam leaving the Q2 collision cell is guided to the TOF pusher region. Since ions of low *m/z* travel faster than heavier ions, some ions will be lost depending on the exact timing of the orthogonal ion pusher. Panel (e) shows how, in Zeno SWATH and ZT Scan DIA, (fragment) ions are being ejected from the Quadrupole region in a pulsed fashion using the Zeno trap and IQ3. Ions are sequentially ejected from higher to lower *m/z* so that more ions arrive simultaneously at the pusher region, equalizing (*m/z*-wise) and increasing the relative ion amounts that can be sent to the TOF analyzer region, thus yielding higher sensitivity.

Here we introduce ZT Scan DIA, an acquisition method that couples scanning quadrupole ion isolation to ion trapping in the Zeno Trap (Fig. 1c,e). Compared to Zeno SWATH it has conceptual benefits. First, continuous quadrupole scanning avoids the overhead of emptying the Q2 collision cell of fragment ions that negatively impacts the duty cycle when decreasing the cycle time (i.e., < 80% at 6.6 ms per MS/MS). Given that Zeno SWATH tends to display optimum performance with acquisition schemes that utilize large numbers of narrow isolation windows and short accumulation times, eliminating this overhead in ZT Scan DIA is particularly beneficial. Second, utilizing a scanning quadrupole with adjustable Q1 isolation width comes with a gain in specificity for assigning fragment ions to peptide precursor ions that surpasses common DIA methods and even approaches the specificity of data-dependent acquisition (DDA or IDA) mode. At the same time, the Zeno Trap efficiently prevents ion losses commonly observed with Scanning SWATH, leading to a higher sensitivity for ZT Scan DIA. Benchmarking ZT Scan DIA as implemented on a QTOF 7600+ prototype instrument (SCIEX), we show that it substantially increases sensitivity, especially in fast proteomic experiments with low sample amounts, and that it improves quantitative precision in complex matrices.

## 2 Materials and methods

### Reagents

LiChrosolv LC-MS grade water and sample preparation reagents were purchased from Merck (Merck, Darmstadt, Germany), acetonitrile from VWR (Darmstadt, Germany), and formic acid from Thermo Fisher Scientific (Waltham, MA, USA). The purchased protein digest standards were K562 Protein Extract Digest (Sciex), MS Compatible Yeast Protein Extract (Promega), and MassPREP *E. coli* Digest Standard (Waters). Working stocks were prepared according to the manufacturer’s instructions (typically in 0.1% formic acid in LC-MS grade water), aliquoted, and stored at –80°C until further use. Multi-species mixes were prepared freshly by mixing thawed and well-mixed stock solutions before diluting to 250 or 50 ng/µl and then being aliquoted into HPLC glass vials.

### Preparation of cell line proteome digest

Human embryonic kidney 293 cells (“HEK”) were cultivated adherently in Dulbecco’s Modified Eagle’s Medium + 10% fetal calf serum (Gibco) + 1% Penicillin / 5% Streptomycin under 5% CO_2_ in an incubator (Bender) heated to 37°C. Upon cell harvest using a cell scraper, they were processed using a urea-based in-solution digestion procedure. In brief, cells were harvested, and 7 M Urea + 100 mM ammonium bicarbonate buffer was added. Cells were homogenized using a GenoGrinder bead beater (SPEX), cleared by centrifugation, and proteins quantified using the BCA assay (Thermo). Upon reduction with 5 mM dithiothreitol for 1.5 h at 30°C and alkylation with 10 mM iodoacetamide for 30 min at room temperature in the dark, Trypsin (Promega) was added in a 100:1 protein-to-protease ratio followed by incubation overnight at 37°C. Next, the digests were quenched by adding formic acid to about 1% (v/v). For the peptide clean-up, we used 96-well C18 solid-phase extraction plates (The Nest Group) following the manufacturer’s protocol. Eluted peptides were dried using a vacuum centrifuge (Eppendorf) and stored at -80°C until needed. Dried peptides were resuspended in 0.1% (v/v) formic acid and aliquoted into glass HPLC vials (Waters).

### LC-MS

#### Overall Instrumental Setup

We used a prototype Zeno trap-enabled Sciex QTOF, similar to the ZenoTOF 7600+ system (Sciex), operated under a development version of Sciex OS 3.

For capillary flow-rate chromatography, the mass spectrometer was interfaced with an OptiFlow source with the SteadySpray Low Micro Electrode (Sciex) connected to a Waters ACQUITY M-Class UPLC (Waters) equipped with a 150 × 0.3 mm Kinetex 2.6 µm XB-C18 column (Phenomenex) that was heated to 30°C while running at 5 µl per minute.

For analytical flow-rate chromatography, we connected the mass spectrometer to a 1290 Infinity II UHPLC system (Agilent). The 5.2-min active gradient measurements employed an Infinity Lab Poroshell 120 EC-C18 2.1 × 50 mm 1.9 µm column (Agilent). For the 3.1-min active gradient experiments, we used a 2.1 × 30 mm Luna OMEGA 1.6 μm C18 100 Å column (Phenomenex). Chromatographic buffers were water + 0.1% (v/v) formic acid as buffer A or acetonitrile + 0.1% (v/v) formic acid as buffer B. Columns were kept at 30°C during analysis.

Analyte ionisation leveraged the OptiFlow source (Sciex) for capillary flow and the Turbo V source (Sciex) for analytical flow rate chromatography.

#### Liquid Chromatography Gradients

**Tab. 1:** Chromatographic Gradient - capillary flow - 15 min active / 27 min total.

|  |  |  |  |  |  |  |  |  |  |  |  |  |  |  |
| --- | --- | --- | --- | --- | --- | --- | --- | --- | --- | --- | --- | --- | --- | --- |
| [min] | 0 | 1 | 2.5 | 5 | 8 | 11 | 13 | 15 | 16 | 16.5 | 17 | 19 | 20 | 27 |
| [%B] | 3 | 5 | 9.5 | 15 | 19 | 22 | 24 | 27 | 32 | 40 | 80 | 80 | 3 | 3 |
| [μl/min] | 5 | 5 | 5 | 5 | 5 | 5 | 5 | 5 | 5 | 5 | 5 | 5 | 5 | 5 |

**Tab. 2:** Chromatographic Gradient - capillary flow - 7 min active / 15 min total.

|  |  |  |  |  |  |  |  |  |
| --- | --- | --- | --- | --- | --- | --- | --- | --- |
| [min] | 0 | 1 | 6 | 6.5 | 7 | 9 | 10 | 15 |
| [%B] | 3 | 3 | 30 | 40 | 80 | 80 | 3 | 3 |
| [μl/min] | 5 | 5 | 5 | 5 | 5 | 5 | 5 | 5 |

**Tab. 3:** Chromatographic Gradient - capillary flow - 3 min active / 15 min total.

|  |  |  |  |  |  |  |  |
| --- | --- | --- | --- | --- | --- | --- | --- |
| [min] | 0 | 1 | 4 | 5 | 7 | 8 | 15 |
| [%B] | 3 | 3 | 30 | 80 | 80 | 3 | 3 |
| [μl/min] | 5 | 5 | 5 | 5 | 5 | 5 | 5 |

**Tab. 4:** Chromatographic Gradient - analytical flow - 5.2 min active / 7.5 min total.

|  |  |  |  |  |  |  |  |
| --- | --- | --- | --- | --- | --- | --- | --- |
| [min] | 0 | 5 | 5.2 | 5.21 | 5.4 | 5.5 | 7.5 |
| [%B] | 3 | 36 | 80 | 80 | 80 | 3 | 3 |
| [μl/min] | 800 | 800 | 800 | 1200 | 1200 | 1000 | 1000 |

**Tab. 5:** Chromatographic Gradient - analytical flow - 3.1 min active / 6 min total.

|  |  |  |  |  |  |  |  |
| --- | --- | --- | --- | --- | --- | --- | --- |
| [min] | 0 | 3 | 3.1 | 4.5 | 4.51 | 5.95 | 6 |
| [%B] | 3 | 36 | 80 | 80 | 3 | 3 | 3 |
| [μl/min] | 800 | 800 | 1200 | 1200 | 1200 | 1200 | 800 |

#### Mass Spectrometry - Zeno SWATH DIA

In each method using capillary flow rate chromatography, we used 12 psi ion source gas 1, 60 psi ion source gas 2, 35 psi curtain gas, 4500 V source voltage, and a source temperature of 300 °C. The MS1 scan range covered *m/z* 400-900 and used an accumulation time of 50 ms, while for MS2, we scanned from *m/z* 140-1750 using 13 or 6.7 ms MS2 accumulation times (see **Tab. 6**) using 65 variable m/z windows for precursor isolation (see **Tab. 7**). Signal intensities were derived from summing signals over 8 time bins.

**Tab. 6:** Zeno SWATH DIA Method Scan speeds per capillary flow rate LC gradients.

| Active Gradient time [min] | MS2 accumulation times [ms] | Cycle time [s] |
| --- | --- | --- |
| 15 | 13 | 1.233 |
| 5 | 6.7 | 0.769 |
| 3 | 6.7 | 0.769 |

<u>Tab. 7 & 8: Zeno SWATH - DIA variable windows and collision energies</u>

For the experiments using analytical flow rate chromatography, ion source gases 1 & 2 and the curtain gas were set to 60, 65, and 55 psi, respectively. The CAD gas was set to 7 psi, the source temperature to 600°C, and the spray voltage to 4000 V. The TOF MS covered *m/z* 400-1500 using 30 ms accumulation time. Precursor ions falling between *m/z* 400 and 900 were isolated using 80 variable windows (see **Tab. 8**) and their fragments detected between *m/z* 100 and 1500 while using 11 ms or 13 ms MS2 accumulation times (see **Tab. 9**). Signal intensities were derived from summing signals over 8 time bins. Collision energies (*CE*) were calibrated depending on the m/z range of interest using the following formula: CE 0.05 * m/z + 5

<u>Tab. 7 & 8: Zeno SWATH - DIA variable windows and collision energies</u>

**Tab. 9:**
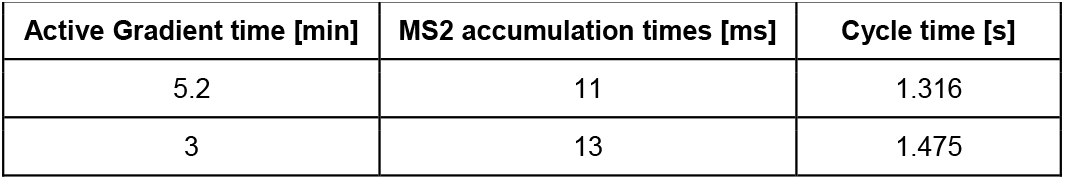
Zeno SWATH DIA Method Scan speeds per analytical flow rate LC gradients.

#### Mass Spectrometry - ZT Scan DIA

Our source region parameters matched the ones described above, in dependence of the applied flow regime. We used three predefined mass spectrometric methods for ZT Scan DIA, available in Sciex OS 3.4 upon purchase of a dedicated license, that account for the different peak widths in response to narrower or wider peptide peaks – 375 Da/s and 7.5 Da Q1 for peaks more or equal to 4.5 s profile-width-at-half-height (PWHH) (230 Hz), 750 Da/s and 10 Da Q1 for peak widths between 1.1 and 4.5 s PWHH (320 Hz), and 750 Da/s and 5 Da Q1 for peak widths less than or equal to 1 second PWHH (641 Hz) (also see **Tab. 11**). The applied collision energies increased with *m/z* adjusted to tryptic peptide precursors at charge state +2. Within Sciex OS 3.4, the acquired ZT Scan DIA raw data undergoes calibration on residual precursor signals and is subsequently converted to SCIEX’s proprietary .wiff- and .wiff.scan-file format, promoting compatibility with commonly used proteomics data processing software tools. In our hands, the obtained file sizes ranged between 1 to about 14 GB, depending on method length and acquisition speed.

**Tab. 11:** Additional ZT Scan DIA Method Acquisition Metrics.

| ZT Scan DIA method | Dwell times [ms] | Cycle time [s] |
| --- | --- | --- |
| 5 Da Q1 - 750 Da/s | 6.7 | 0.779 |
| 10 Da Q1 - 750 Da/s | 13.3 | 0.786 |
| 7.5 Da Q1 - 375 Da/s | 20 | 1.459 |

### Data processing

The mass spectrometric raw data were processed using DIA-NN 1.8.1 (Demichev et al., 2020). For measurements involving only human proteins, we used the same spectral library as in Wang et al. (Wang et al., 2022) which was generated from high pH reverse-phase fractionated K562 and HeLa human proteome digests in IDA/DDA on a capillary flow LC - ZenoTOF 7600 MS system that got searched with ProteinPilot within the OneOmics suite against a human Swiss-Prot canonical and isoform database (UniProt Consortium, 2025). The results were concatenated using the Extractor application in the OneOmics suite (SCIEX) to create the spectral library. For the LFQ bench-type analysis (Navarro et al., 2016), we started from an *in-silico* predicted spectral library of *Homo sapiens, Saccharomyces cerevisiae*, and *Escherichia coli* including only SwissProt curated proteins including isoforms (downloaded in July 2023).

MS2 mass accuracies were allowed to fall within a 20 ppm tolerance while those for MS1 were inferred automatically (i.e., set to “0”). The scan window was set to 6 for micro-flow and 7 for analytical-flow chromatography acquisitions. For method performance evaluations, the match-between-runs (MBR) feature was enabled for replicates using the same LC-MS settings (e.g., replicates within the parameter screening and LFQ mixes of a given LC-MS setup using the same sample amount). Library generation was set to “Smart profiling” and “Heuristic protein inference” was enabled for all measurements. The FDR threshold was set to 1% on the precursor level per run for all analyses.

Data post-processing and analysis were performed using Python 3.9 with the pandas 2.2.3, numpy 2.1.3, and scipy 1.14.1 packages. DIA-NN reports were filtered depending on the experimental goal: Precursor-centric ID benchmarks were filtered to equal or less than 1% FDR on Q.Value (run-specific Precursor FDR) and Lib.Q.Value. Likewise, Protein-centric benchmarks were filtered to less than or equal to 1% FDR on PG.Q.Value and Lib.PG.Q.Value. Coefficients of variation (CVs) were obtained by dividing the standard deviation of signal intensities with a degree of freedom N = n - 1 by its mean. CVs were only accepted for identifications with n ≥ 3 and are indicated as values or percentages. LOESS regression was performed within the Seaborn package using the regplot function under default settings. All plots were generated in Python using the Seaborn 0.13.2 package and arranged using Inkscape 1.2 and 1.4.

## 3 Results

### Combining the Zeno Trap and Scanning SWATH to ZT Scan DIA

In Zeno trap-enabled scanning data-independent acquisition (“ZT Scan DIA”), ions move through the instrument in a similar manner as presented previously (Messner et al 2021) (Fig.1c,e). Likewise, the Q1 quadrupole scans the precursor ion space with a set Q1 isolation window and speed. To maintain sufficient points across the chromatographic peak, it is important to couple the Q1 speed to the expected peak width during chromatographic separation. Given the current status of chromatography, peak widths are generally about 4.5 s or less across the majority of different chromatographic setups. To remove the need for fine-tuning methods related to different peak width at half height (PWHH), SCIEX released three optimized methods (> 4.5 / 1 < x ≤ 4.5 / ≤ 1 seconds, respectively, corresponding to 375 Da/s with 7.5 Da Q1, 750 Da/s with 10 Da Q1, and 750 Da/s with 5 Da Q1, see Methods and Fig. S1), enabling users to select the best method tailored to their experimental needs. Precursor ions transmitted by the scanning Q1 reach the Q2 quadrupole stage where collisional ion activation and fragmentation occur. The acquisition software automatically selects collision energies and an MS/MS fragment mass range suitable for bottom-up proteomics workflows. Next, the generated fragment ions are collected via the Zeno trap, which releases them in pulses at a 1500 Hz frequency to the orthogonal TOF pulser. This pulsed release from the Zeno trap increases the TOF duty cycle, enhancing ion coverage, particularly for low m/z ions (Loboda & Chernushevich, 2009). To maximize the benefit of the Zeno trap, the Q1 scanning speeds were set to align with the Zeno trap pulsing frequency, ensuring optimal dwell times for fragment ion signal collection in terms of signal-to-noise and points-per-peak ratios. The acquisition software organizes ion detection events into specific bins based on the TOF pulses that overlap with the precursor m/z, allowing for the extraction of precursor profiles for any given fragment (Fig. 1c, right). This results in tightly correlating fragment ion signals and precursor m/z, allowing for the differentiation of true analyte signals from noise with greater specificity.

To systematically compare ZT Scan DIA with Zeno SWATH DIA, we used microflow rate chromatography (5 µl/min on M-Class, Waters) connected to a prototype Zeno trap-enabled QTOF 7600 system (SCIEX) and matched method parameters like the 400-900 *m/z* range and MS/MS accumulation times of Zeno SWATH DIA with dwell times of ZT Scan DIA, meaning we spent a comparable time per fragment ion signal collection. We measured a human K562 cell line proteome tryptic digest standard (SCIEX) with 15- and 7-minute active separation times, respectively. The obtained data were subsequently processed using DIA-NN with a spectral library obtained from data-dependent acquisitions on a deep-fractionated K562 digest (Methods).

First, we focused on repeated injections of the standard sample while monitoring the consistency of identification. Injecting 5 ng of K562 digest, corresponding to a proteome mass from about 20-30 mammalian cells (Kulak et al., 2014), we quantified on average 14,769 and 16,950 precursors from 2,586 and 3,017 proteins with a median CV of 19.4% and 16.5% on Protein-level for Zeno SWATH and ZT Scan DIA, respectively, using a 15-minute active gradient. Using the faster 7-minute gradient, we obtained averages of 1,991 and 2,440 proteins (from 12,165 and 14,861 precursors), respectively, with median CVs of 17.0% and 19.0% (Fig. S2). This corresponds to an increase of +15% on unique Precursors and +17% Protein groups for 15-minute active gradients, and +22% and +23% for 7-minute active gradients on average for ZT Scan DIA, compared to Zeno SWATH DIA, respectively (Fig. 2a,d). Moreover, in combination with the 15-minute active gradient, ZT Scan DIA consistently identifies more proteins at the same data completeness level while for the 7-min gradient, higher data completeness appeared for Protein groups that were detected in less than about 80% of the replicates of each method (Fig. 2b,e). Inspecting the quantitative precision for shared precursors across intensity quantiles yielded an overall improved precision for 15-minute gradients, and a comparable precision to Zeno SWATH DIA with 7-minute active gradients (Fig. 2c,f).

**Fig. 2:**
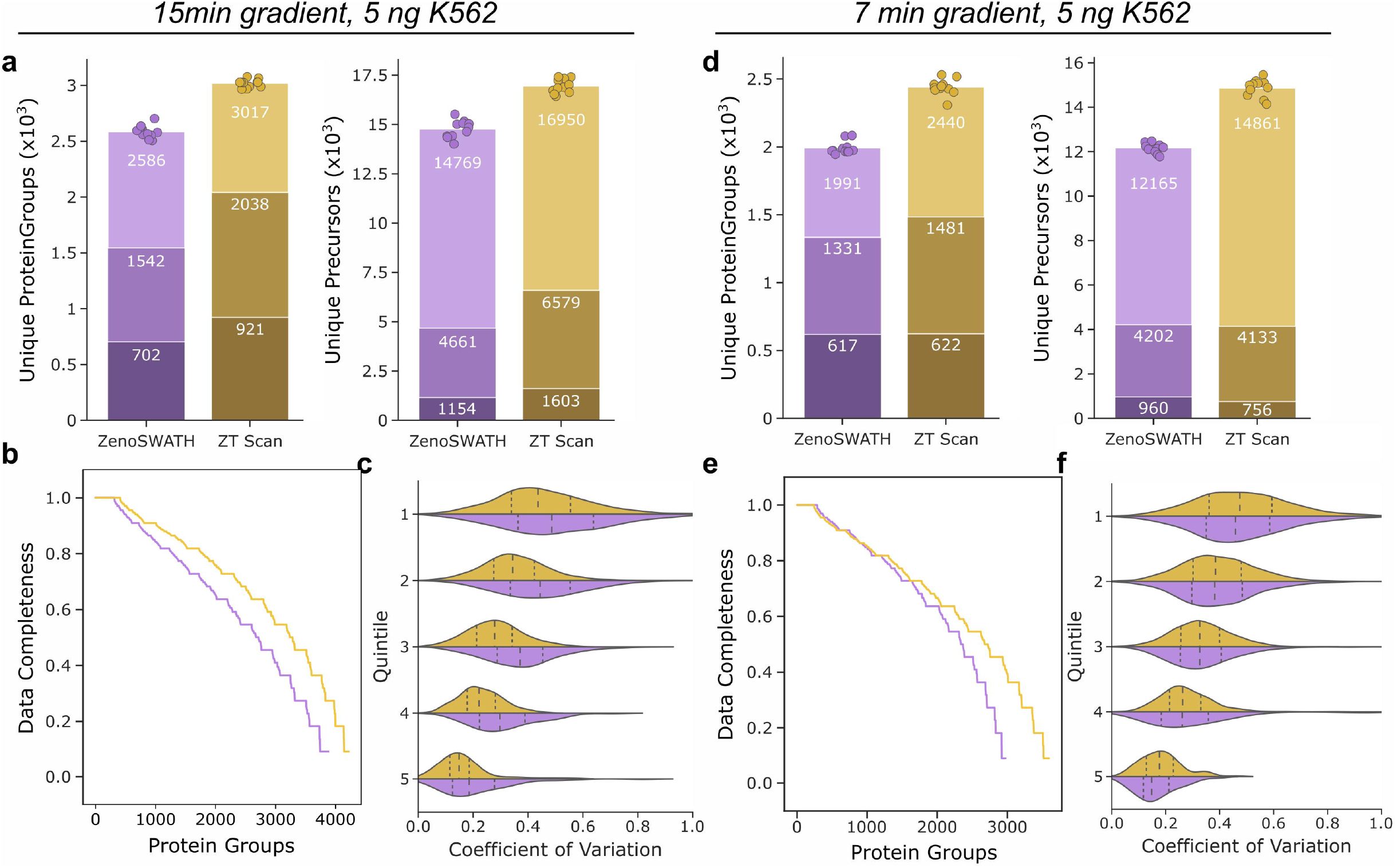
Performance Assessment of ZT Scan DIA compared to Zeno SWATH DIA on replica injections. K562 human proteome digest standard was acquired in eleven replicates of 5 ng using 15 and 7-minute active gradients. Panels (a) and (d) highlight the number of unique Protein groups or Precursors at 1% FDR per replicate injection. Panel (a): ZT Scan DIA predominantly detects more precursors and protein groups on average, with better than 20% CV and below 10% CV (bars stacked top to bottom) than Zeno SWATH DIA. Panels (b) and (e) highlight improved data completeness for ZT Scan DIA across the replicate series (same color code as in upper panels). Panels (c) and (f) showcase the achieved quantitative precision for jointly detected precursors for both methods at different flow regimes across the replicate series. Quintiles were derived from splitting the intensity range into five similarly sized chunks. ZT Scan DIA yields higher precision for 15 min gradients and comparable performance when using a 7 min gradient.

Next, we expanded our analysis by probing different sample loads at different acquisition speeds. We tested 200, 50, 5, and 1 ng of K562 using the same chromatographic gradients (15- and 7-min active gradients) and 50 and 200 ng K562 digest on a fast 3-minute active gradient, respectively. To account for narrower peaks during faster chromatography, we adjusted each ZT Scan DIA method according to the chromatographic peak widths (as above, Methods) regarding their dwell times compared to Zeno SWATH DIA MS2 accumulation times. Using the 15-minute active separation gradient and 200 ng K562, we observed comparable identification numbers between Zeno SWATH and ZT Scan DIA, respectively, but ZT Scan DIA had a slightly better quantitative precision. Moreover, it yielded more precisely quantified proteins (i.e., below 20% CV) when reducing the load to 50, then 5, and 1 ng of the K562 digest. Notably, the lower the injection amounts, the higher the gain of ZT Scan DIA compared to Zeno SWATH DIA. For example, for the lowest load of 1 ng, ZT Scan DIA quantified 42% more protein groups (Fig. 3a). On the shorter gradients, 7-min and 3-min (Fig. 3b,c), ZT Scan DIA resulted in more overall and also more quantitatively confident identifications (Fig. 3d-f), especially with the 7-min active gradient separation (Fig. 3g-i).

**Fig. 3:**
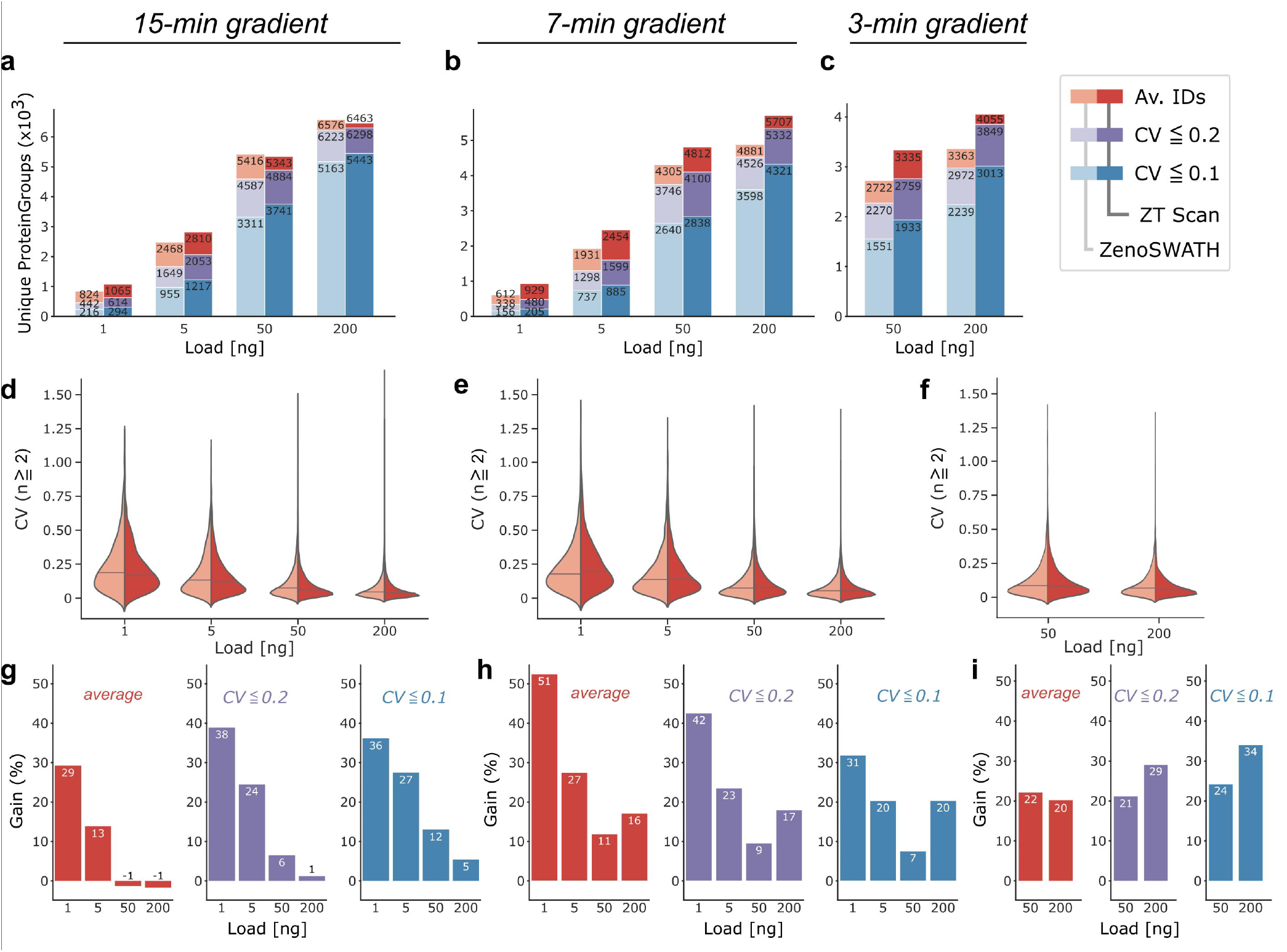
Identification Performance Assessment of ZT Scan DIA compared to Zeno SWATH DIA on Titration Series. K562 human proteome digest standard was acquired using 15-min and 7-min gradients at 200, 50, 5 and 1 ng loadings, and 200 and 50 ng at 3-min gradients. Panels (a-c) highlight the number of unique Protein groups at 1% FDR from triplicate injections with changing sample amounts loaded (average: red, _≤_20% CV: purple, _≤_10% CV: blue; Zeno SWATH and ZT Scan DIA shown with lower and higher saturation, respectively). ZT Scan DIA generally detects more protein groups and at better precision. The less sample was loaded, the better ZT Scan DIA fared compared to Zeno SWATH DIA. Panels (d-f) confirm comparable overall quantitative precision if not lower CVs when using ZT Scan DIA (dark red) compared to Zeno SWATH DIA (light red). Medians are indicated with a line. The bottom panels (g-i) highlight the gains in IDs per load across the quality categories as in the panels above (same colour code as in the top row).

### Quantitative Performance of ZT Scan DIA

Next, we used multi-species proteome deconvolution (“LFQ-bench-type”) experiments (Navarro et al., 2016) to assess the quantification performance of the new hybrid method. These assays benchmark the quantitative performance of a proteomic method through their ability to deconvolute proteomes of different species mixed at different ratios (Fig. 4a). We measured five mixes composed of varying amounts of three proteome standard digests (*Homo sapiens* (K562 cell extract tryptic digests), *Saccharomyces cerevisiae* (whole-cell protein extract), and *Escherichia coli* (purified cytosolic protein fraction)), at two different loads using active gradients corresponding to 15- and 7-min active gradient setups, respectively. The measurements were conducted in triplicate and on the aforementioned microliter-flow rate LC-MS platform. Comparing ZT Scan with Zeno SWATH DIA resulted in ∼6% more protein identification rates for 250 ng injections using Zeno SWATH DIA, while for 50 ng injections, ZT Scan DIA increased protein identification by +38% on average for 15- and 7-min active gradients (Fig. 4b-d). Besides the identification performance of any LC-MS proteomics method, a typical pitfall related to quantitation is ratio compression for peptide precursors in the lower signal intensity quantiles that arise from higher signal-to-noise ratios for these low-abundant analytes. Further, detector noise and interfering signals from co-fragmented peptide ions may lead to a decline in quantitative precision. Both methods performed similarly across all tested conditions when assessing the quantitative precision for protein groups expressed as coefficients of variation (Fig. S3). Concerning quantitative accuracy, however, ZT Scan DIA returned favorably over Zeno SWATH DIA when loading 250 ng and a 15-minute active gradient, i.e., the squared errors of observed to expected ratios were smaller (Fig. 4k & Fig. S4). Likewise, when reducing the sample load or shortening the chromatographic separation time, ZT Scan DIA again provided higher quantitative accuracy. Notably, the quantitative ratios acquired with ZT Scan DIA were more stable across the intensity range for all LC-MS conditions and mixes tested (Fig. 4e-m & Fig. S4).

**Fig. 4:**
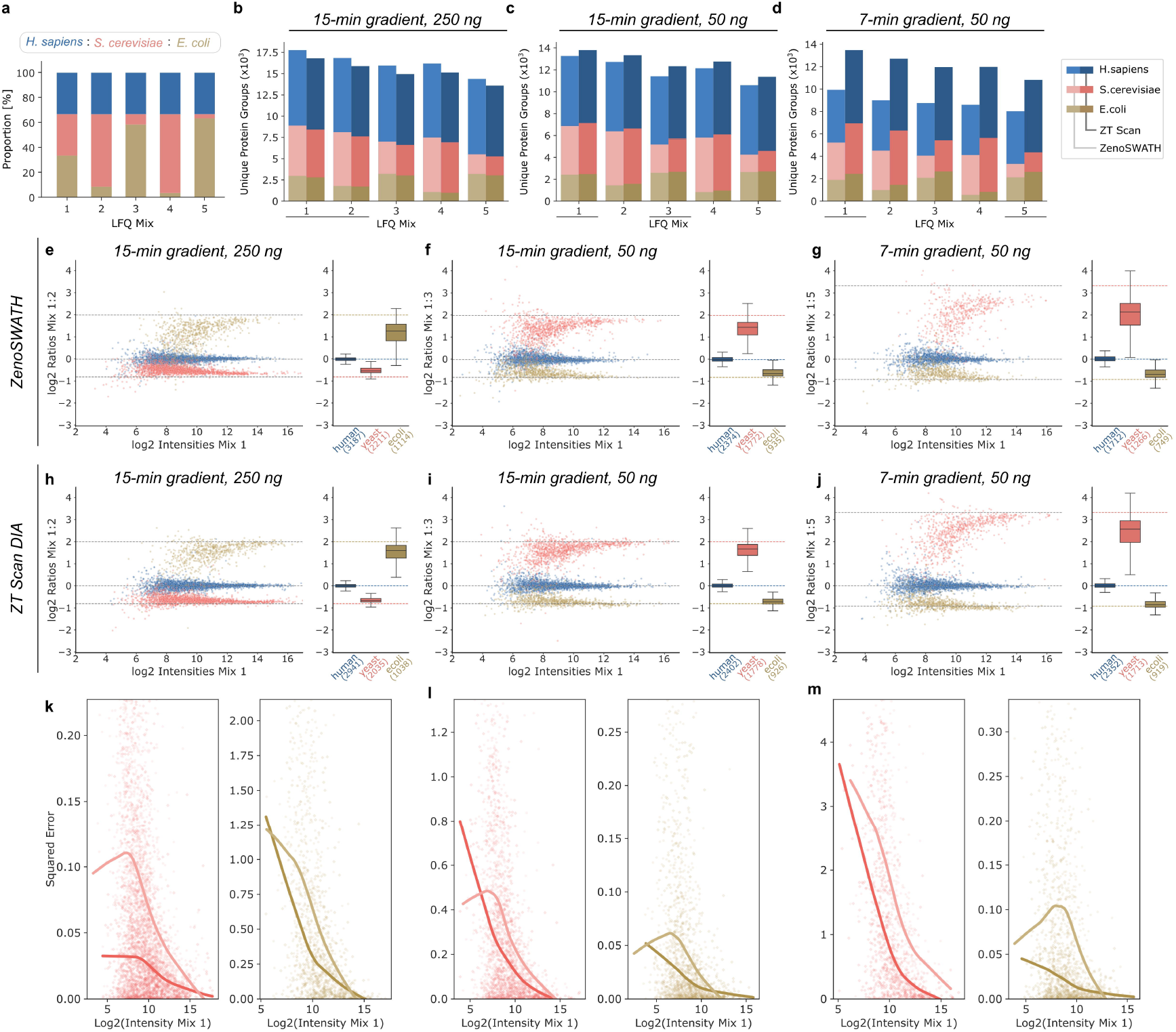
Label-free quantitation benchmark of ZT Scan DIA compared to Zeno SWATH DIA. Multi-species proteome deconvolution (“LFQ-bench type”) analysis using mixes of human, yeast, and bacterial proteome tryptic digests at defined ratios as shown in panel (a) were acquired in triplicate, with three different LC-MS acquisition regimes as indicated on top of panels (b-d). Panels (b-d) show the obtained protein group quantities per mix and by species for active gradients of 15 / 15 / 7 min active gradient at 250 / 50 / 50 ng load, respectively. Panels (e-g) highlight the quantitative performance of Zeno SWATH DIA in selected LFQ-bench experiments. Panels (h-j) show the corresponding LFQ-bench plots using ZT Scan DIA. Data was filtered for Protein groups detected in at least two replicates and the resulting averaged quantities were offset to match a human log ratio median of zero. Panels (k-m) highlight the quantitative error in the accuracy of each protein group quantity on the deviant species for both methods based on the squared error to the expected ratio across the intensity range. Lines represent LOESS regressions. ZT Scan DIA data points are shown as diamonds while Zeno SWATH DIA as points, colored as in panels (b-d) above.

In summary, ZT Scan DIA reduced ratio compression effects for protein quantities at extreme ratios compared to Zeno SWATH DIA, resulting in more accurate protein quantification across the covered detection range.

### Proteomic experiments with analytical flow rate chromatography and ZT Scan DIA

Next, we assessed the performance of ZT Scan DIA employing analytical flow rate chromatography, as a strategy for addressing the need for high-throughput proteomics applications with fast gradients to be run robustly and precisely over large sample series, suited for experiments were maximizing proteomic depth is secondary to throughput and quantification precision (Messner et al., 2021, 2023). The key to maximising the performance of DIA-based proteomics methods in analyzing proteomes with analytical flow rate chromatography is matching data acquisition speeds to its high chromatographic peak capacity (Ludwig et al., 2018; Messner et al., 2021). If the right balance is struck, the attainable analytical specificity and sensitivity will be maximized due to reduced signal interferences from co-occurring peptide signals (i.e., resolved chromatographically and via fast-changing m/z isolation windows) and diminished signal loss per analyte (i.e., from a higher sampling rate). Besides aiding the peptide identification process in DIA proteomics, higher sampling rates cause an increase in the number of points-per-peak, benefiting precise peptide quantitation. For example, under our global acquisition parameters, Q1 scanning at 750 Da/s with an isolation window of 5 Da equates to an MS/MS rate of about 640 Hz for ZT Scan DIA, which compares favorably to achievable TOF MS/MS rates of about 133 Hz using Zeno SWATH DIA. In addition, fragment ions are observed in multiple consecutive scans with ZT Scan DIA instead of only being seen within the next DIA cycle in Zeno SWATH. Thus, in theory, ZT Scan DIA should allow for superior identification rates at excellent quantification reliability in demanding acquisition regimes aiming for ultimate data acquisition speeds.

We tested two chromatographic gradients for analytical flow rate separations, a 5.25-minute and a 3.1-minute active gradient, comparing ZT Scan DIA methods to an optimized Zeno SWATH DIA method while analyzing 0.5 / 1 / 2 µg of an in-house generated human embryonic kidney (HEK) cell line digest (Methods). Comparing results for the 5.25-minute gradient, we observe increasing gains in parallel with higher sample loads culminating in a plus of about 1000 protein groups detected when injecting 1 or 2 µg of the peptide mixture using the ZT Scan DIA method optimized for ca. one-second wide peaks (5 Da Q1 & 750 Da/s Q1 scanning)(+22% / +27%). Accordingly, the numbers of precisely quantified proteins and precursors from 2 µg of sample increased by +31% / 39% for Protein groups and +48% / 61% for Precursors with CVs of 20% / 10%, respectively (Fig. 5a,c). Higher gains were achieved with the 3.1-minute chromatography method, from which we detected 4515 Protein groups on average with 5 Da Q1 & 750 Da/s ZT Scan DIA, representing a gain of +33% over Zeno SWATH DIA. Considering precursors with trustworthy quantities using the 10 Da Q1 & 750 Da/s ZT Scan DIA method, we note that precursors below 10% CV almost outnumber those of Zeno SWATH DIA even when including all below 20% CV (Fig. 5b,d). Consequently, when the focus is on getting the greatest number of precisely quantified precursors (<10% CV), the ZT Scan DIA method for intermediate peak widths (10 Da Q1 & 750 Da/s) fared best while after aggregation to Protein groups, this advantage over the fastest ZT Scan DIA method (5 Da Q1 & 750 Da/s) was offset and accordingly, the latter method may be used (Fig. 5b,d & Fig. S5). Notably, ZT Scan DIA yielded highly precise proteomic measurements, with Protein group CV medians well below 20% (Fig. 5e,f). Regarding identification overlaps, all methods showed highly consistent identification performance within their replicates, but also among them, with a slight tendency for ZT Scan DIA methods to share more identifications (Fig. S6 & S7). Moreover, we compared signal responses between methods and at different sample loads (Fig. 5g-j). Albeit analyte quantities from Zeno SWATH DIA displays high correlation comparing single-shot LC-MS acquisitions of 2 µg and between 2 and 0.5 µg HEK loadings (Pearson Correlation Coefficient (“Rho”) of 0.945 with 19,390 / 0.84 with 14,824 precursor ratios), ZT Scan DIA performed even better with Rho’s of 0.980 and 0.902 from 22,988 and 17,257 precursor quantity ratios, respectively. Thus, despite an increase of 19% in precursor identifications, ZT Scan DIA demonstrates comparable or better signal correlation, again maintaining an edge over Zeno SWATH DIA. Overall, ZT Scan DIA led to a more precise quantitation in a high-throughput proteomics setting.

**Fig. 5:**
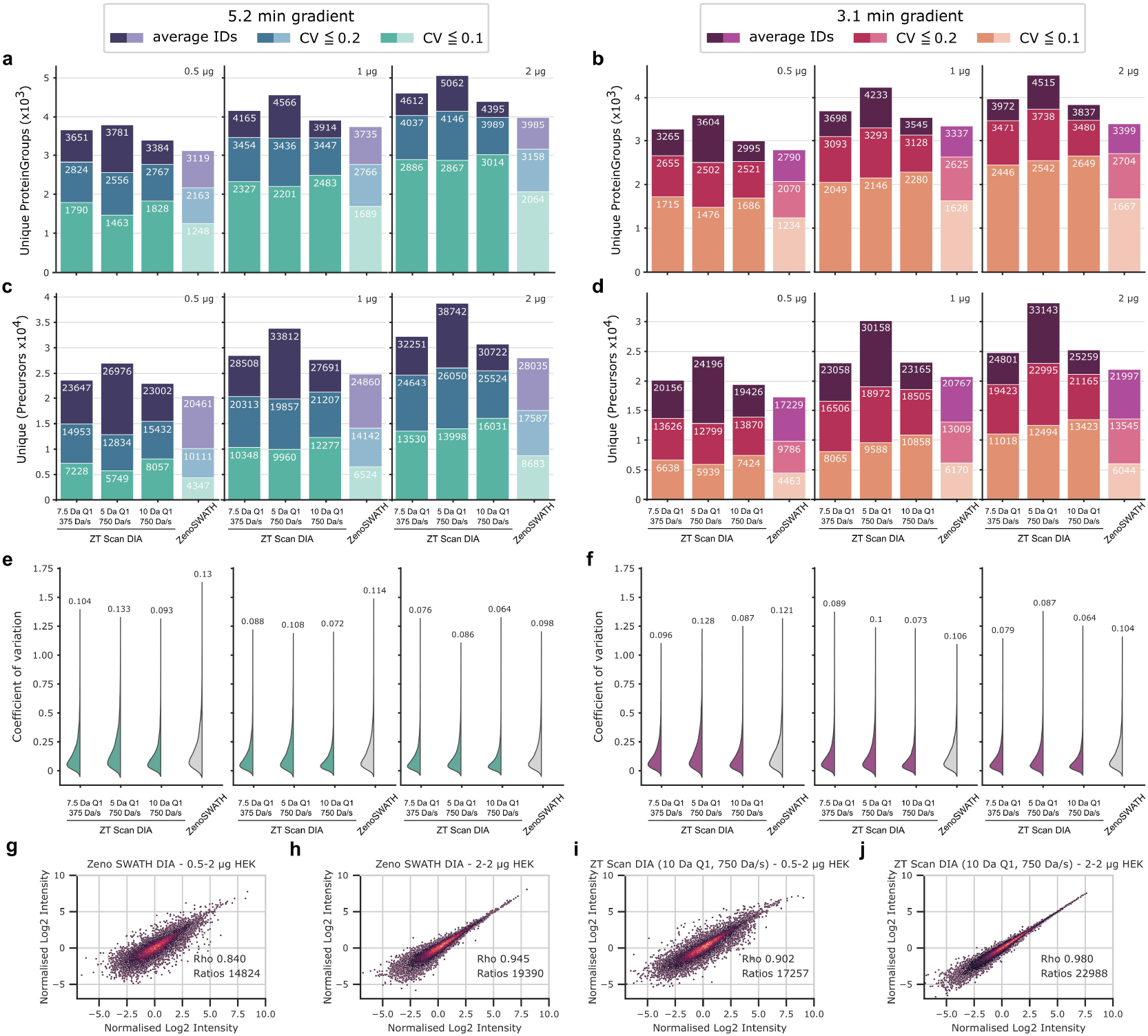
ZT Scan DIA Improves High-Throughput Proteomics on a prototype ZenoTOF 7600+ System. Assessment of ZT Scan DIA’s performance in analytical flow rate LC-MS. Panels (a-d) show identifications of protein groups and precursors at increasing sample loads of 0.5 / 1 / 2 µg human proteome digest using a 5.25- or 3.1-minute active gradient compared to an optimized Zeno SWATH DIA methodology in three categories: average identifications from triplicate, identifications below or equal to a CV of 0.2, and of 0.1 (from top to bottom, n=3), respectively. Numbers for each category are indicated below a bar’s top. Panels (e & f) demonstrate improved quantitative precision for all ZT Scan DIA methods tested over Zeno SWATH DIA. ZT Scan DIA outperformed Zeno SWATH DIA in total identifications and those precisely quantified. Panels (g-j) show signal correlations for different methods and at different sample loads using the 3.1-minute active gradient chromatography as follows: (g) Zeno SWATH DIA, 2 µg each, 2 replicates, (h) Zeno SWATH DIA, 0.5 and 2 µg, 2 replicates, (i) ZT Scan DIA, 0.5 and 2 µg, 2 replicates, and (j) ZT Scan DIA, 2 µg each, 2 replicates. Pearson Correlation coefficients and numbers of valid ratios are indicated. The data was filtered to 1% FDR on the respective level.

## 4 Discussion

While data-independent acquisition methods increasingly dominate in proteomic experiments (Fröhlich et al., 2024; Jiang et al., 2024), there is an ongoing need to increase their performance in terms of acquisition speed, as well as quantitative precision and accuracy. We here propose an acquisition method, ZT Scan DIA, that combines the scanning dimension that we introduced with Scanning SWATH (Messner et al., 2021) with the Zeno SWATH acquisition method (Wang et al., 2022). In ZT Scan DIA, the fast-scanning Q1 quadrupole adds extra specificity over the conventional SWATH approach due to fast-changing staggered precursor ion isolation windows to trace fragment ion correlation with high temporal resolution and which effectively reduces adverse signal interferences while the rapid ion processing in the Zeno trap adds extra sensitivity by increasing the TOF duty cycle.

Implemented on a prototype Zeno TOF 7600+ instrument (SCIEX), we demonstrate ZT Scan DIA’s protein identification and quantification capabilities on several generic proteome standards, benefiting inter-laboratory comparisons. In scenarios with average throughput and low sample inputs (5 ng across eleven replicates), and in comparison to Zeno SWATH DIA on the same setup, ZT Scan DIA increased protein identifications by 22%. Despite the low input material, proteomes were quantified precisely (CVs≤20% for about 2/3rd of all proteins from eleven replicate injections using a 15-minute active gradient). When systematically probing the three different available ZT Scan DIA methods with varying inputs of sample, we found gains for almost all conditions tested with the sweet spot being a 7-minute active gradient, where the inclusion of the scanning dimension increased protein identifications by 23% on average (i.e., across all sample loads and categories - average, CV<20%, CV<10%) and even by 51% for 1 ng input. Thus, ZT Scan DIA is well suited, particularly when only small sample amounts are available or when deeper proteomic coverage at precise quantitation is needed making it an ideal choice for studies on larger cohorts. In addition, in scenarios with abundant sample material, in application with fast analytical-flow rate chromatography using active gradients as fast as 3.1 minutes only is when ZT Scan DIA shines - leading to consistent gains exceeding +30% on protein identifications, while notably also giving improved quantitative precision for up to 40% more proteins when compared to Zeno SWATH DIA.

In summary, we demonstrate the introduction of a sliding Q1 Quadrupole with the Zeno trap enabled QTOF and benchmark its performance in combination with capillary- and analytical flow rate chromatography. We report an increase in sensitivity and quantitative precision at different sample throughput regimes. In the future, we envision ZT Scan DIA’s best applications in these proteomics fields that benefit most from increased specificity and sensitivity. These entail but are not limited to i) high-throughput clinical proteomics which relies on high-precision protein quantification to overcome low effect sizes and other confounders for efficient patient stratification and biomarker identification by allowing robust statistics (Cai et al., 2023; Geyer et al., 2024), ii) proteomics experiments with very low input materials, which are challenged predominantly by a lack of sensitivity and a larger extent of missing values (Geiger, 2023; Rosenberger et al., 2023), as well as iii) system biology studies, such as functional proteomic experiments, time series analysis, or the analysis of strain collections, that all require high sample numbers and are sensitive to batch effects and depend on precise proteome quantification (Messner et al., 2023).

## 5 Associated Data

The mass spectrometric data and DIA-NN logs and reports have been deposited on ProteomeXchange via the PRIDE partner repository (Perez-Riverol et al., 2019) with the dataset identifier https://www.ebi.ac.uk/pride/archive/projects/PXD063462.

## Supporting information

Tables 7&8 - Collision Energy Settings

## Acknowledgments

We acknowledge the Charité Core Facility High Throughput Mass Spectrometry, especially Michael Mülleder, for scientific advice and technical support, and Nir Cohen for proofreading the draft version. This work was supported by the Ministry of Education and Research (BMBF), as part of the National Research Node ‘Mass Spectrometry in Systems Medicine’ (MSCoreSys), under grant agreements 031L0220 (to M.R.) and 161L0221 (to V.D.), and the Sonderforschungsbereich (SFB) TRR 186 (to Z.W.). Work in M.R.’s lab is further supported by the European Research Council (ERC-SyG-2020951475). C.B.M. is supported by the Precision Proteomics Center Davos, which receives funding from the Swiss canton of Grisons.

## Author Contributions

S.T., C.A., C.B.M., L.R.S., and M.R. designed the experiments; D.L. cultivated HEK cells and prepared in-house proteome digests. L.R.S. and Z.W. collected LC-MS data. C.A., A.J., I.B., P.P., and S.T. contributed to technical software and hardware development and data interpretation; L.R.S. wrote the draft; all authors contributed to the manuscript text; J.C.-P., S.T., V.D., and M.R. supervised the study.

