## Supplementary Figures for "Performance Characteristics of Zeno Trap Scanning DIA for Sensitive and Quantitative Proteomics at High Throughput"

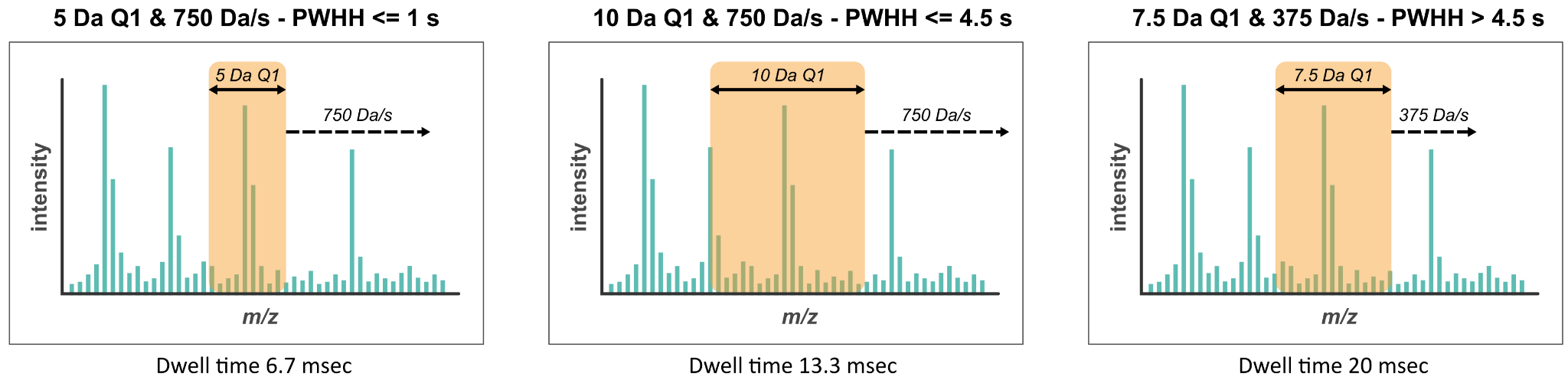

#### Fig. S1: Scheme on scanning DIA precursor isolation for three ZT Scan DIA methods

The user can choose from either of the three ZT Scan DIA methods that differ in their Q1 isolation width (orange boxes) and Q1 scan speeds (dashed arrow) to arrive at individual performance optima depending on a peptide’s chromatographic peak width at half height (PWHH).

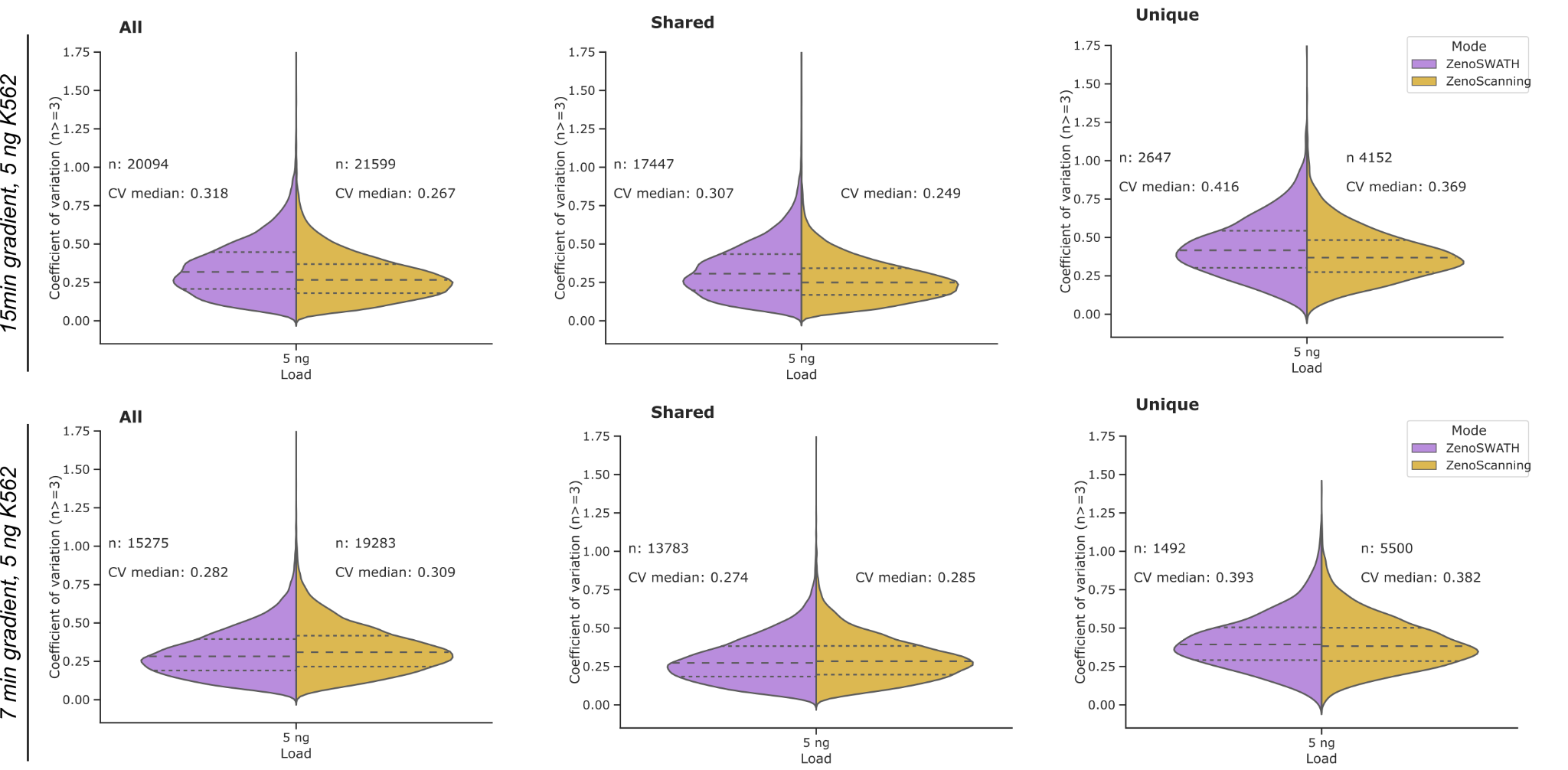

#### Fig. S2: Quantitative precision of precursor from eleven low-load acquisitions comparing ZT Scan DIA with Zeno SWATH DIA

Each panel illustrates the distribution of coefficients of variation for quantified precursors (observed at least three times) per method, in set categories of either all precursors detected per method, being shared between methods, or being unique to each method. Numbers of observations and medians are indicated per method and across categories. The color code is indicated in the right-most panels.

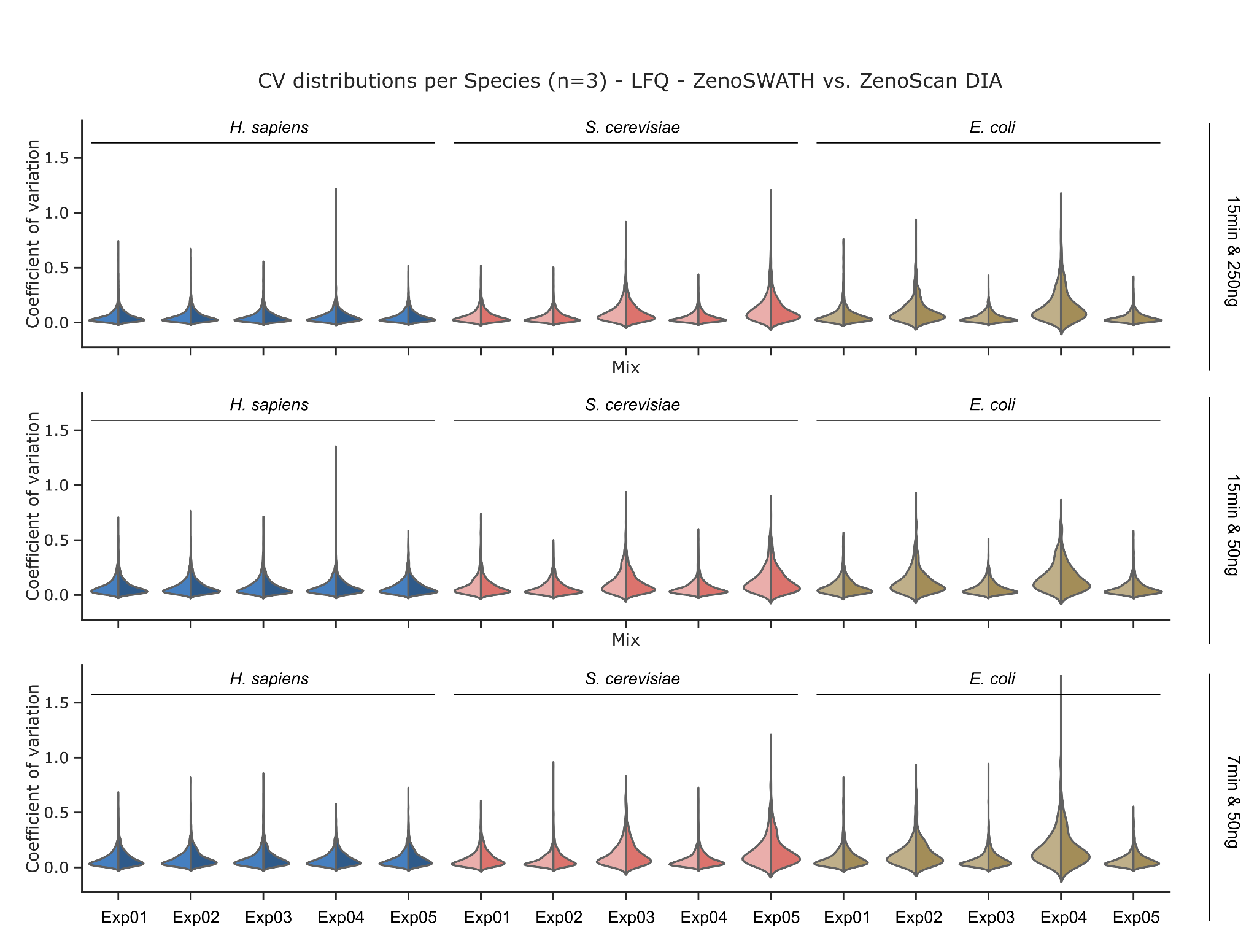

#### Fig. S3: Quantitative precision for precursor identifications in LFQ bench experiment comparing ZT Scan DIA with Zeno SWATH DIA

Violin plots illustrate the distribution of coefficients of variation for precursors observed at least three times, for each experimental mix (refer to Fig. 4a for exact ratios), and per species (human: blue, yeast: red, bacteria: brown). As indicated on the right side, each panel row represents an active gradient (15 or 7 min) and the total amount of loaded sample per LC-MS acquisition (250 or 50 ng).

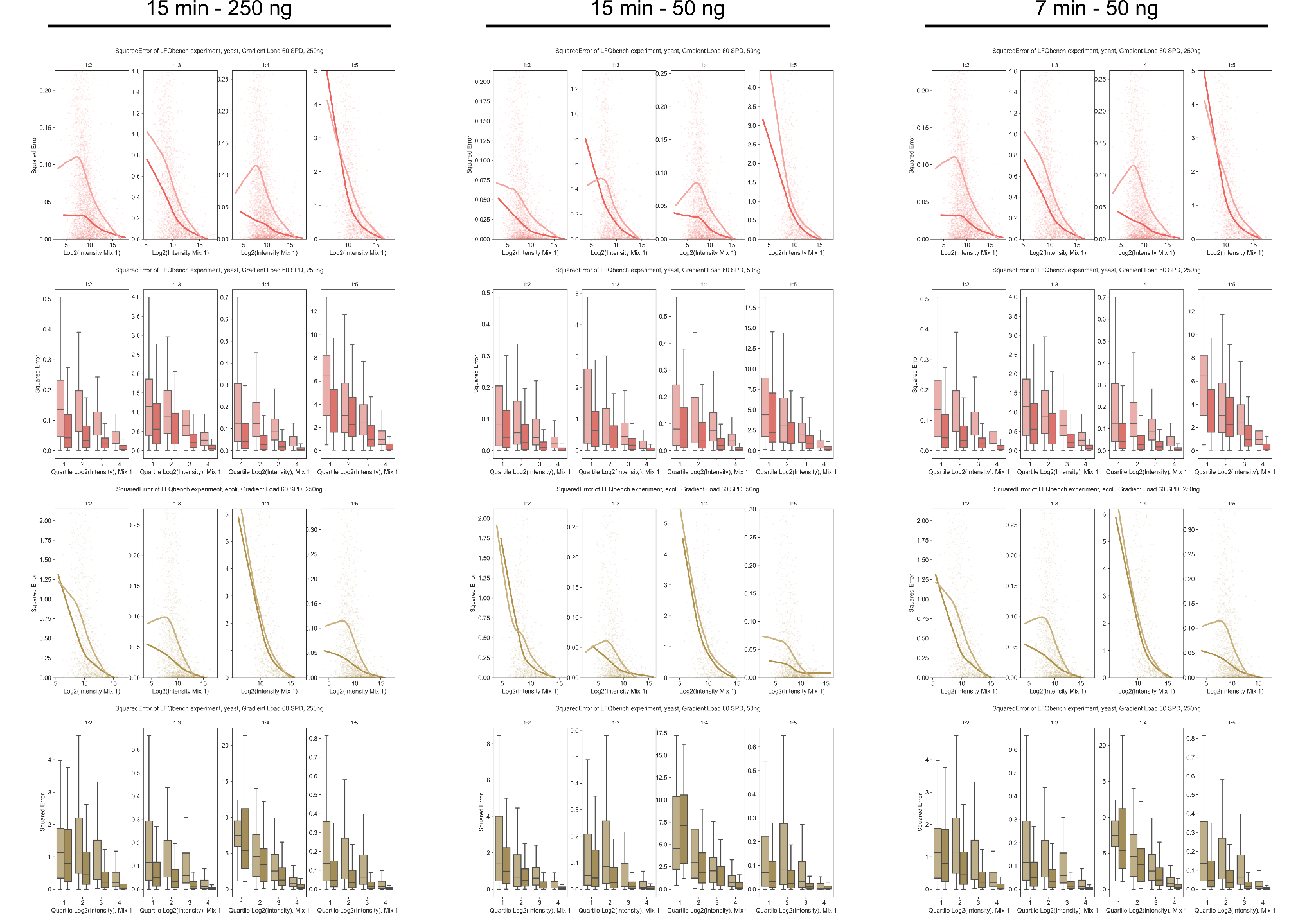

#### Fig. S4: Quantitative accuracy for protein group identifications in LFQ-bench experiment comparing ZT Scan DIA with Zeno SWATH DIA

The quantitative accuracy was assessed based on the difference (as squared error) of a measured quantity to its expected quantity (given the defined mixing ratios). Each column illustrates the experimental condition applied with regard to active chromatographic gradient and total sample load. In each column, each row consists of four panels that indicate the quantitative accuracy for each of the four experimental mixes compared to the standard mix (i.e., to experimental mix 1, with equal relative ratios of all tested proteomes). Datapoints in scatterplots were fitted using LOESS regression to highlight the trends. Boxplots show the same trends across log2 intensity quartiles (boxes cover 50% of the data while the whiskers span 1.5x the IQR; outliers are not shown). Darker colouring highlights the performance of ZT Scan DIA while the lighter hue stands for the corresponding one for Zeno SWATH DIA.

Red colouring indicates errors of yeast protein groups while brown colouring indicates those of *E. coli*.

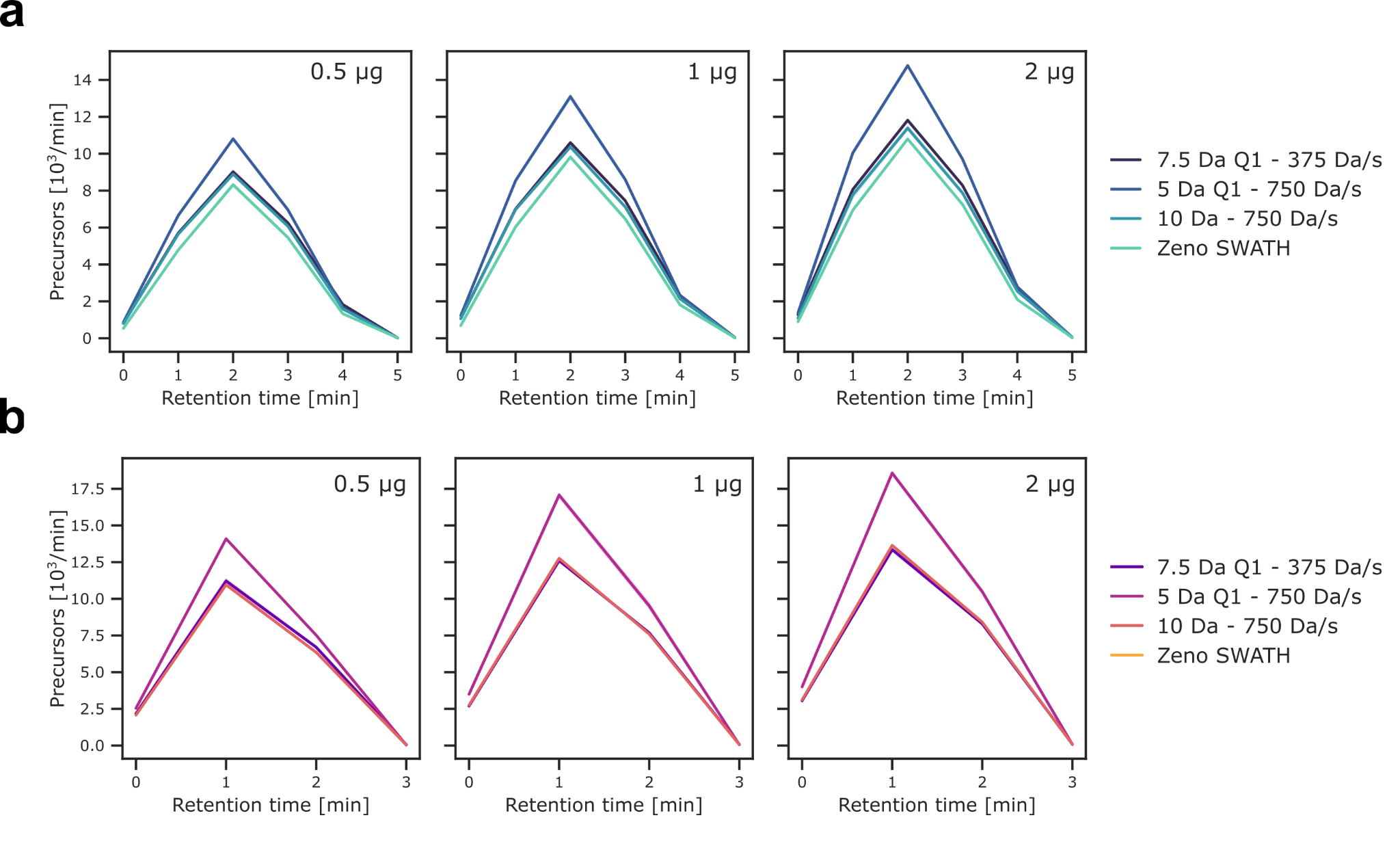

#### Fig. S5: Precursors per minute in analytical flow rate chromatography experiment

Precursors per minute are plotted for each sample load of HEK cell proteome digest, pooled for triplicate injections, for the 5.25-min active gradient (a) and the 3.1-min active gradient (b).

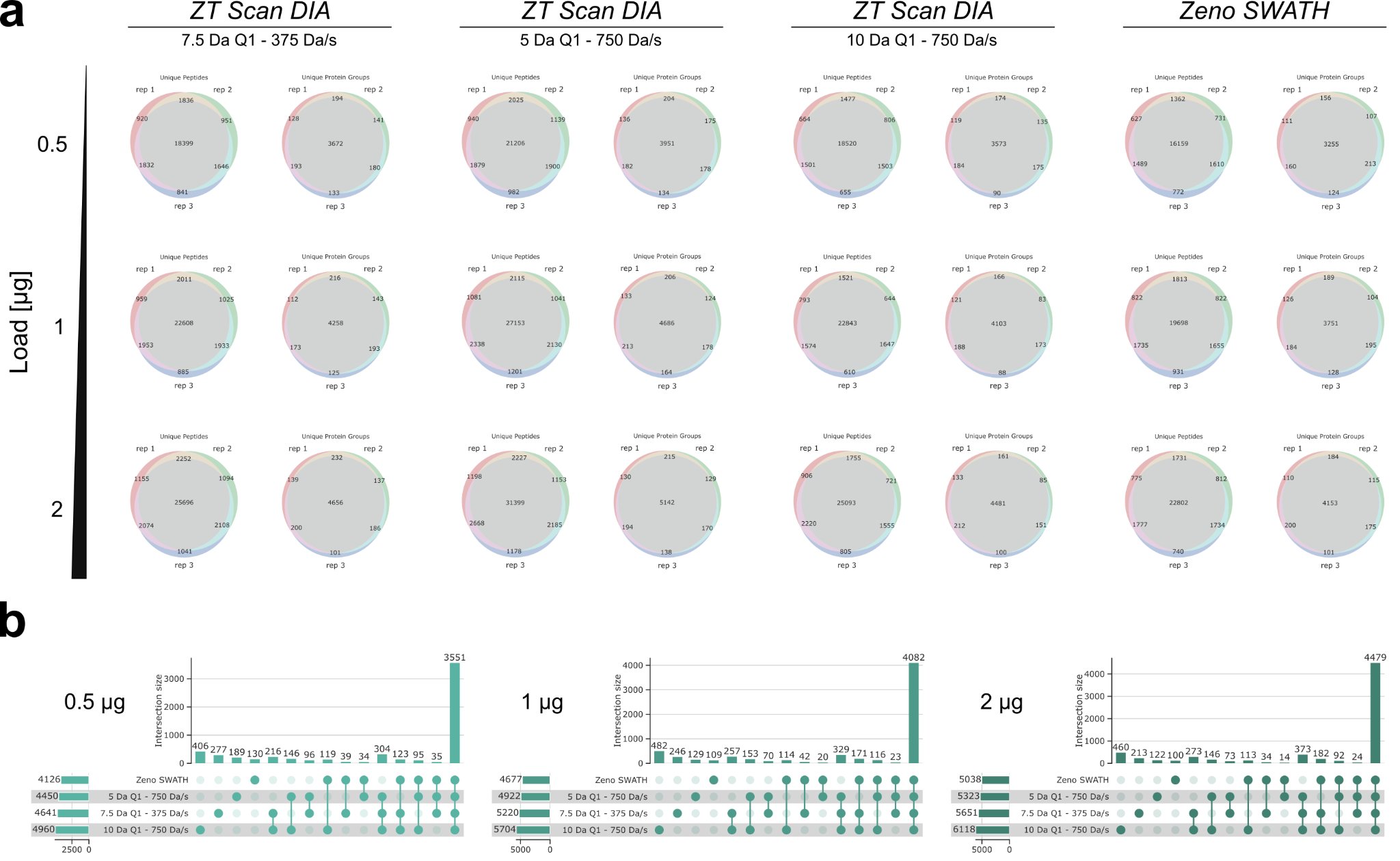

#### Fig. S6: Set overlaps between replicates and both MS methods for 5.2 min active analytical gradient

a) Venn diagrams showcasing overlaps of Peptides (left) or Protein groups (right) within replicates of a loading amount and LC-MS method combination (indicated above). b) Upset Plot illustrating the overlaps of peptide identifications between different LC-MS methods (3x ZT Scan DIA and Zeno SWATH) and sample loading amounts (per panel), pooled for replicates.

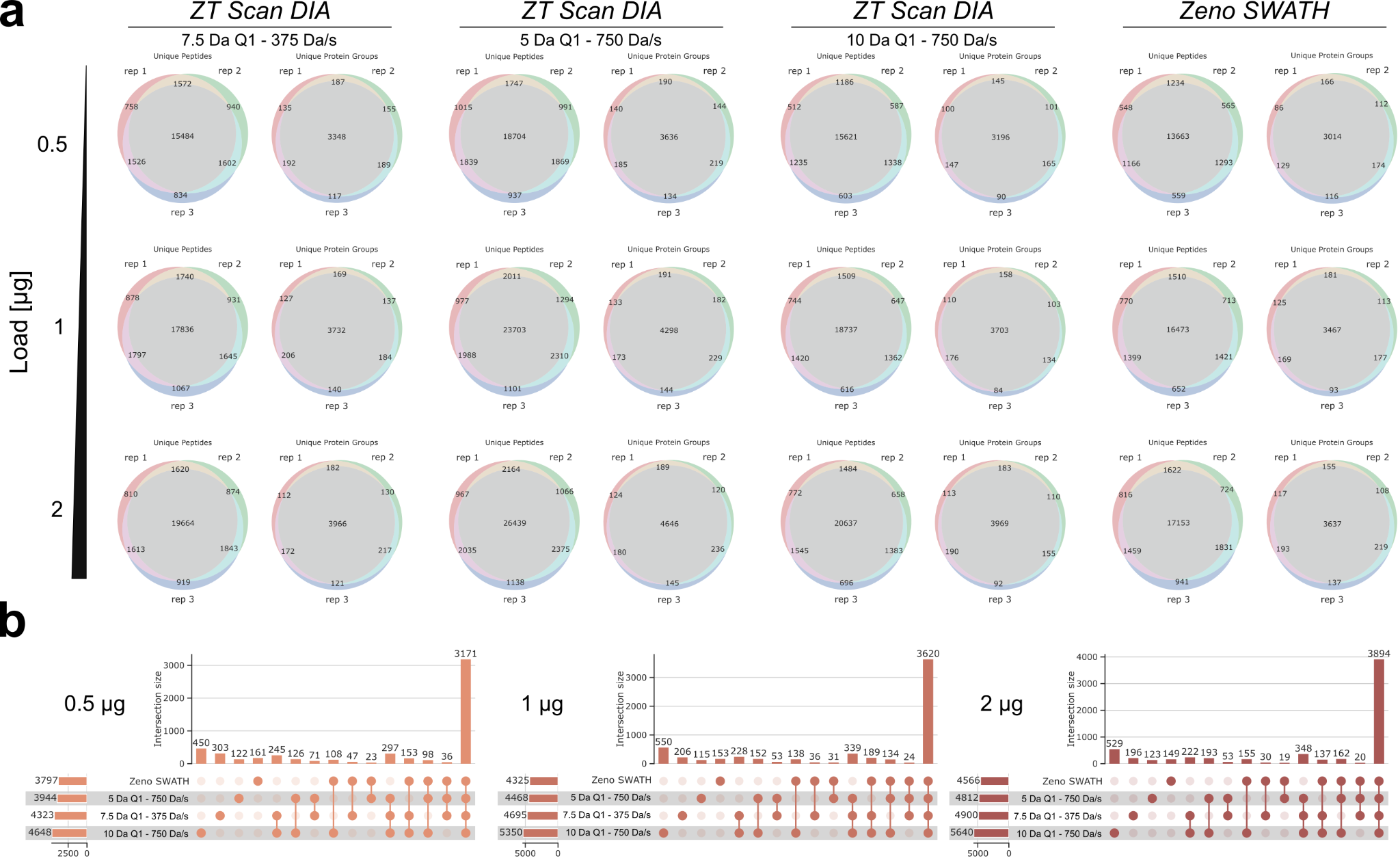

#### Fig. S7: Set overlaps between replicates and both MS methods for 3.1 min active analytical gradient

a) Venn diagrams showcasing overlaps of Peptides (left) or Protein groups (right) within replicates of a loading amount and LC-MS method combination (indicated above). b) Upset Plot illustrating the overlaps of peptide identifications between different LC-MS methods (3x ZT Scan DIA and Zeno SWATH) and sample loading amounts (per panel), pooled for replicates.
